# Host metabolic reprogramming in response to SARS-Cov-2 infection

**DOI:** 10.1101/2020.08.02.232645

**Authors:** S T R Moolamalla, Ruchi Chauhan, U Deva Priyakumar, P K Vinod

## Abstract

Understanding the pathogenesis of SARS-CoV-2 is important for developing effective treatment strategies. Viruses hijack the host metabolism to redirect the resources for their replication and survival. How SARS-CoV-2 influences the host metabolism is still unclear. In this study, we analyzed transcriptomic data obtained from different human respiratory cell lines and patient samples (Swab, PBMC, lung biopsy, BALF) to understand the metabolic alterations in response to SARS-CoV-2 infection. For this purpose, the expression pattern of metabolic genes in the human genome-scale metabolic network model Recon3D was explored. We identified metabolic genes and pathways and reporter metabolites under each SARS-CoV-2-infected condition and compared them to identify common and unique changes in the metabolism. Our analysis revealed host-dependent dysregulation of glycolysis, mitochondrial metabolism, amino acid metabolism, glutathione metabolism, polyamine synthesis, and lipid metabolism. We observed different metabolic changes that are pro- and antiviral in nature. We generated hypotheses on how antiviral metabolism can be targeted/enhanced for reducing viral titers. These warrant further exploration with more samples and in vitro studies to test predictions.

## Introduction

The ongoing pandemic of coronavirus disease (COVID-19) is a highly infectious respiratory illness in humans caused by a strain called Severe Acute Respiratory Syndrome Coronavirus 2 (SARS‑CoV‑2), which is believed to have zoonotic origins(Benvenuto et al., 2020; Zhou et al., 2020). It was first identified in December 2019 in Wuhan and quickly caused more than 14.5 million infections, including at least 606,000 deaths spanning 188 countries by 20th July 2020. WHO has declared this ongoing pandemic as a current global public health emergency. There is no treatment currently available, and disease management includes only critical medical care and social distancing to avoid disease spread. Most patients have one or more symptoms of fever, cough, shortness of breath, headache, lack of taste/smell, chest pain, and diarrhoea, while in severe cases patients might develop severe pneumonia, pulmonary edema, ARDS, or multiple organ failure and lead to death (Chen et al., 2020). The severity is attributed to dysregulation of immune function induced by the virus and host factors like comorbidities; however, their relative contributions are still unknown. It is critical to understand the pathogenesis of SARS-CoV-2 for the design of effective treatment strategies.

Viruses’ life cycle involves entry into a host, evading the host cell immune response, and viral replication by taking control of host-cellular machinery for protein synthesis. These are mediated by the host and virus protein-protein interaction network. The viral mechanism for infecting hosts may vary depending upon the viral type. SARS‑CoV‑2 virus entry into a host human cell depends on the binding of spike protein (S) to the receptor ACE2 and priming of S protein by serine protease TMPRSS2 (Hoffmann et al., 2020). Serine protein inhibitor against TMPRSS2 is shown to block the entry into host cells partially. Further, inhibitors based on the crystal structure of the main protease (M^pro^, 3CL^pro^) of SARS-CoV-2 has been proposed (Dai et al., 2020; Mengist et al., 2020). In addition to viral proteins, different host proteins can serve as targets. Virus-host interactome based drug repurposing strategies have shown great promise in identifying different targets in the host (Cheng et al., 2018; Deisy Morselli Gysi, 2020). However, there is minimal knowledge of the mechanism of the SARS-CoV-2 survival strategy.

Due to their complete dependence on host cells to replicate, viruses may have evolved different strategies to reprogram the host metabolism for their replication and survival (Mayer et al., 2019; Moreno-Altamirano et al., 2019; Sanchez and Lagunoff, 2015; Thaker et al., 2019). Specific host metabolic pathways, including carbohydrate, fatty acid, and nucleotide metabolism, are known to be breached by different viruses upon infection. Each viral species is likely to induce unique metabolic reprogramming of the host cell. The host transcriptomic data in response to SARS-CoV-2 is helping to decipher the changes at the level of gene expression. Elevated inflammatory cytokine production and low innate antiviral defenses in different hosts is shown in response to SAR-CoV-2 infection (Blanco-Melo et al., 2020; Lieberman et al., 2020; Xiong et al., 2020). Autophagic and mitochondrial dysfunction have been reported with specific cell lines (Singh et al., 2020). However, a comprehensive understanding of metabolic reprogramming of the host is still lacking for SARS-CoV-2.

In this study, we explored the metabolic alterations induced by the SARS‑CoV‑2 virus in different human respiratory cell lines and samples collected from patients. We identified metabolic hot-spots and their corresponding regulatory elements under each condition. A comparison across multiple conditions helped to identify common and unique changes in the metabolism. This analysis generates insights into host metabolic response to SARS-CoV-2, which can be targeted for the effective antiviral response.

## Methods

We used RNA sequencing (RNASeq) raw read count data obtained from 4 human cell lines: adenocarcinomic alveolar basal epithelial (A549), ACE2 transduced A549 (A549-ACE2), human adenocarcinomic lung epithelial (Calu3) and bronchial epithelial (NHBE) infected with SARS-CoV-2 (**Table 1**) (Blanco-Melo et al., 2020). A549 and A549-ACE2 cell lines were infected with a SARS-CoV-2 viral load, multiplicity of infection (MOI) equal to 0.2 and 2, whereas Calu3 and NHBE were infected with a viral load, MOI=2. We also used RNASeq raw read count data obtained from lung biopsy (LB) (Blanco-Melo et al., 2020) and nasopharyngeal swabs of healthy and SARS-CoV-2 infected human samples (Lieberman et al., 2020). We performed differential gene expression analysis using the DESeq2 (v1.26.1) in R (v 3.6.1) for each dataset to obtain differentially expressed genes (DEGs) of SAR-CoV-2 (Love et al., 2014). We also obtained DEGs for bronchoalveolar lavage fluid (BALF) and peripheral blood mononuclear cells (PBMC) cells of SARS-CoV-2 patients reported by Xinog et al., (2020). We compared the DEGs across conditions and identified key genes based on the number of occurrences. The KEGG pathways associated with DEGs were obtained using EnrichR (adjusted p-value < 0.05) (Kuleshov et al., 2016). In order to specifically identify metabolic gene alterations, we filtered the resulting DEGs based on the metabolic genes present in a comprehensive genome-scale metabolic network model, Recon3D (Brunk et al., 2018).

**Table 1:** Transcriptomics data used to study the metabolic response of host to SARS-CoV-2.

| Accession No. | Cell Lines/<br>Regions | Viral Load<br>MOI | Description | Platform |
| --- | --- | --- | --- | --- |
| **GSE147507**<br>**(GEO)** | A549 | 0.2 | Human Lung adenocarcinomic alveolar basal epithelial cells | Illumina NextSeq 500 |
|  | A549 | 2 | Human Lung adenocarcinomic alveolar basal epithelial cells |  |
|  | A549-ACE2 | 0.2 | A549 cells transduced with a vector expressing human ACE2 |  |
|  | A549-ACE2 | 2 | A549 cells transduced with a vector expressing human ACE2 |  |
|  | Calu3 | 2 | Human adenocarcinomic lung epithelial cells |  |
|  | Lung Biopsy | N/A | Lung biopsy from human postmortem |  |
|  | NHBE | 2 | Primary human bronchial epithelial cells |  |
| **PRJCA002326**<br>**(BioProject)** | BALF | N/A | Bronchoalveolar lavage fluid | Illumina HiSeq 2000 |
|  | PBMC | N/A | Peripheral blood mononuclear cells | MGISEQ-2000 and Illumina NovaSeq |
| **GSE152075**<br>**(GEO)** | Swabs | N/A | Human Nasopharyngeal Swabs | Illumina NextSeq 500 |

The DEGs of SARS-CoV-2 were used to identify the metabolites around which the most significant transcriptional changes occur, known as reporter metabolites (Patil and Nielsen, 2005). We used the Recon3D model to identify the neighboring genes of each metabolite. The Recon3D model contains 10,600 metabolic reactions, 5,835 metabolites, and 1,882 unique metabolic genes spanning nine compartments. The p-values (p_i_) obtained from the differential gene expression analysis were transformed into the Z-scores (Z_i_) using the inverse normal cumulative distribution function (CDF). Metabolites were assigned a Z-score (Z_metabolite_) by aggregating the Z scores of their ‘k’ neighboring genes. These Z_metabolite_ scores were corrected for the background distribution using mean (μ_k_) and standard deviation (σ_k_) of aggregated Z scores obtained by sampling 10000 sets of k enzymes from the network. Corrected Z-scores were then transformed to p-values by using CDF and the metabolites with p-values less than 0.05 were identified as reporter metabolites.

We used the recent host-viral protein-protein interactome of SARS-CoV-2 to identify the metabolic DEGs targeted by viral proteins (Gordon et al., 2020). In addition to direct interactions, we also retrieved the metabolic DEGs targeted indirectly by viral targets in the immediate neighborhood of DEGs. To extract the immediate neighbors of host metabolic proteins, we used high confidence and curated human protein-protein interaction network hPPiN (Sambarey et al., 2017). Networks were visualized in Cytoscape version 3.3. Further, we identified the transcription factors associated with differentially expressed metabolic genes. The binding motifs from the promoter of these genes were scanned to obtain significantly over‐represented transcription factor binding site motifs (p-value < 0.05). We performed the binding site analysis in Enrichr using the TRUST transcription factors 2019 database (Kuleshov et al., 2016) for each condition and studied the overlap.

## Results and Discussion

We identified DEGs under each condition and compiled a compendium of all the changes that can serve as a useful resource on SARS-CoV-2 (**Table S1)**. This information can be used to extract consensus and condition-specific genes. Genes of cytokine-mediated signaling pathway are present in most conditions representing the adverse inflammatory condition (“cytokine storm”) of SARS-CoV-2. In this study, we used this resource to specifically identify metabolic alterations in various cell lines and human samples infected by SARS-CoV-2. We extracted the genes of the human genome-scale metabolic model, Recon3D, that are differentially expressed between normal and infected conditions (p-value < 0.05) (**Table S2**). In general, A549 and Calu3 have the most metabolic changes compared to other cell lines/tissues. The overlap of metabolic DEGs across different cell lines/tissues is shown in **Figure 1**. We observe that most metabolic DEGs are downregulated in A549_ACE2, A549, Calu3, lung biopsy and swab compared to NHBE, BALF, and PBMC samples. The metabolic DEGs of lung biopsy, BALF, swab, and PBMC show overlap with the cell line data. However, we found less DEG overlap between these samples obtained from SARS-CoV-2 infected patients. Further, A549_ACE2 shows some metabolic overlap with the A549 cell line. There is also an increase in the number of DEGs with an increase in viral load from 0.2 to 2. Swab samples have very few DEGs compared to other conditions, and most DEGs are downregulated. **Table 2** shows the top metabolic candidate genes upregulated in most conditions. We used the metabolic DEGs to perform the pathway enrichment analysis and identify reporter metabolites.

**Table 2:** Top metabolic genes (p < 0.05) upregulated in response to SARS-CoV-2. Number of occurrences of each gene in different host conditions are given. A549 and A549-ACE2 is shown for MOI of 0.2 (A549_0.2, ACE2_0.2) and 2 (A549_2, ACE2_2).

| Gene | Occurrences | Samples |
| --- | --- | --- |
| NAMPT | 8 | A549_0.2, A549_2, ACE2_0.2, ACE2_2, Calu3, Lung Biopsy, NHBE, Swabs |
| SAT1 | 8 | A549_0.2, A549_2, ACE2_0.2, ACE2_2, Calu3, Lung Biopsy, NHBE, Swabs |
| SOD2 | 7 | A549_0.2, A549_2, ACE2_0.2, ACE2_2, Calu3, Lung Biopsy, NHBE |
| ASS1 | 6 | A549_0.2, A549_2, ACE2_0.2, Calu3, NHBE, BALF |
| CMPK2 | 6 | A549_0.2, ACE2_0.2, ACE2_2, Calu3, Lung Biopsy, Swabs |
| PDE4B | 6 | A549_0.2, ACE2_0.2, ACE2_2, Calu3, Lung Biopsy, Swabs |
| PTGS2 | 6 | A549_0.2, A549_2, ACE2_0.2, ACE2_2, Calu3, NHBE |
| ABCA1 | 5 | A549_2, ACE2_2, Calu3, BALF, Swabs |
| B4GALT1 | 5 | A549_2, ACE2_0.2, ACE2_2, Calu3, NHBE |
| CSGALNACT1 | 5 | A549_0.2, A549_2, ACE2_2, Calu3, NHBE |
| DUOX2 | 5 | A549_2, ACE2_2, Calu3, NHBE, BALF |
| EXT1 | 5 | A549_0.2, A549_2, ACE2_2, Calu3, BALF |
| GCH1 | 5 | A549_2, ACE2_0.2, ACE2_2, Calu3, Lung Biopsy |
| GPCPD1 | 5 | A549_0.2, A549_2, ACE2_0.2, ACE2_2, Calu3 |
| IREB2 | 5 | A549_0.2, A549_2, ACE2_0.2, ACE2_2, Calu3 |
| KYNU | 5 | A549_0.2, A549_2, ACE2_2, Calu3, NHBE |
| LAP3 | 5 | A549_0.2, ACE2_2, Calu3, NHBE, PBMC |
| NT5E | 5 | A549_0.2, A549_2, ACE2_2, Calu3, BALF |
| PTEN | 5 | A549_2, ACE2_0.2, ACE2_2, Calu3, BALF |
| SLC7A2 | 5 | A549_0.2, A549_2, ACE2_0.2, ACE2_2, Calu3 |
| SLC9A8 | 5 | A549_2, ACE2_0.2, ACE2_2, Calu3, Lung Biopsy |
| TYMP | 5 | A549_0.2, A549_2, ACE2_2, Calu3, NHBE |

**Figure 1:**
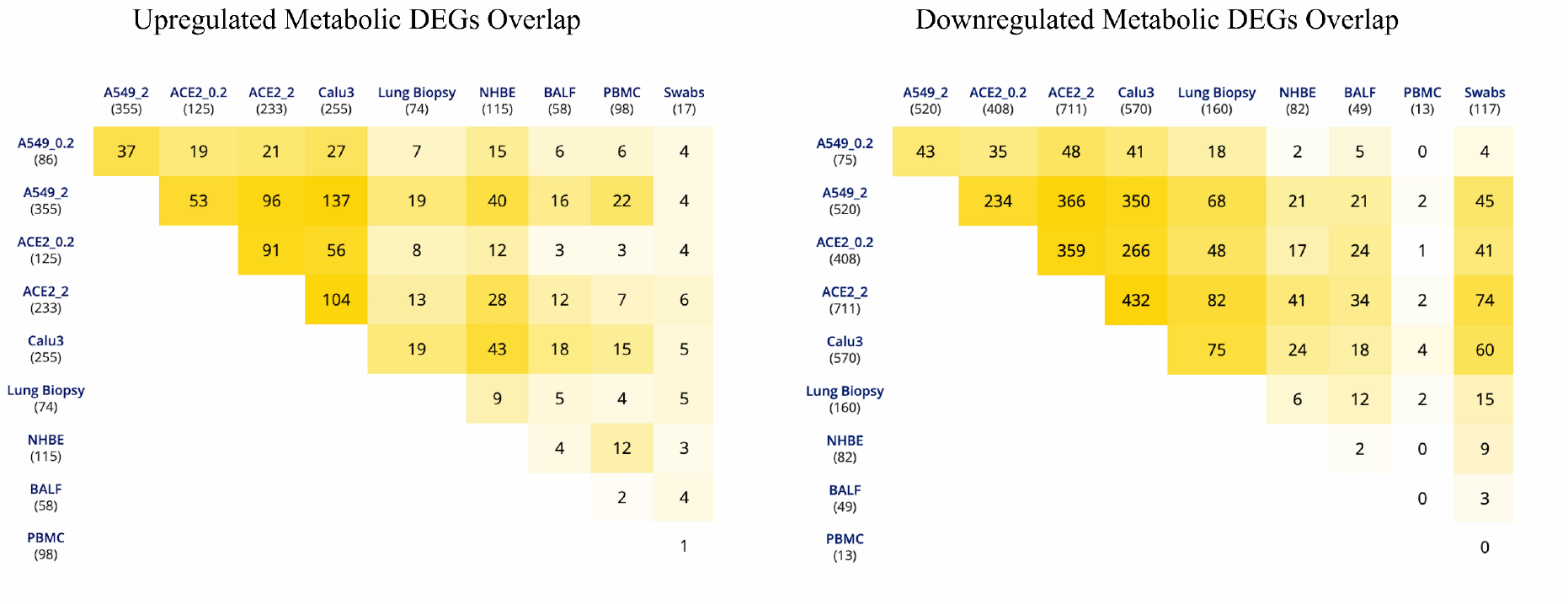
Overlap of differentially expressed metabolic genes in different cell lines and patient samples with SARS-CoV-2 infection.

### Carbohydrate metabolism

The enrichment of downregulated metabolic DEGs show downregulation of tricarboxylic acid cycle (TCA cycle) and oxidative phosphorylation in most cell lines (**Figure 2**). A549_ACE2 cell lines show mitochondrial dysfunction even at a low viral load (ACE2_0.2). This suggests that SARS-CoV-2 does not rely on mitochondrial metabolism for replication and survival. Mitochondria is shown to have a role in host innate immunity and can be a target for invading microorganisms. The mitochondrial antiviral mechanism involves the activation of retinoic acid-inducible gene I (RIG-I)-like receptors (RLR) and activation of its target mitochondrial antiviral (MAV) at the mitochondrial outer membrane (MOM). MAV recruits effectors at the MOM to form MAV signalosome upon viral infection and control the activation of NFκB and IRF3 (Refolo et al., 2020; Seth et al., 2005). The inhibition of MAV results in increased viral replication and the killing of host cells. The antiviral innate immunity is governed by the mitochondrial membrane potential (Δψ_m_) (Koshiba, 2013). RLR antiviral signaling also depends on oxidative phosphorylation (Yoshizumi et al., 2017). Further, we also found downregulated genes are associated with neurodegenerative disorders and non-alcoholic fatty liver disease (NAFLD). This is also related to mitochondrial dysfunction in neurodegenerative disorders, which can activate the NLRP3 inflammasome pathway (Duan et al., 2020). Therefore, mitochondria dysfunction can lead to sustained inflammasome signaling in SARS-Cov-2 (Holley and Schroder, 2020). On the other hand, the TCA cycle and oxidative phosphorylation genes are upregulated in PBMC. Mitochondrial metabolites are upregulated in PBMC (**Figure 3**).

**Figure 2:**
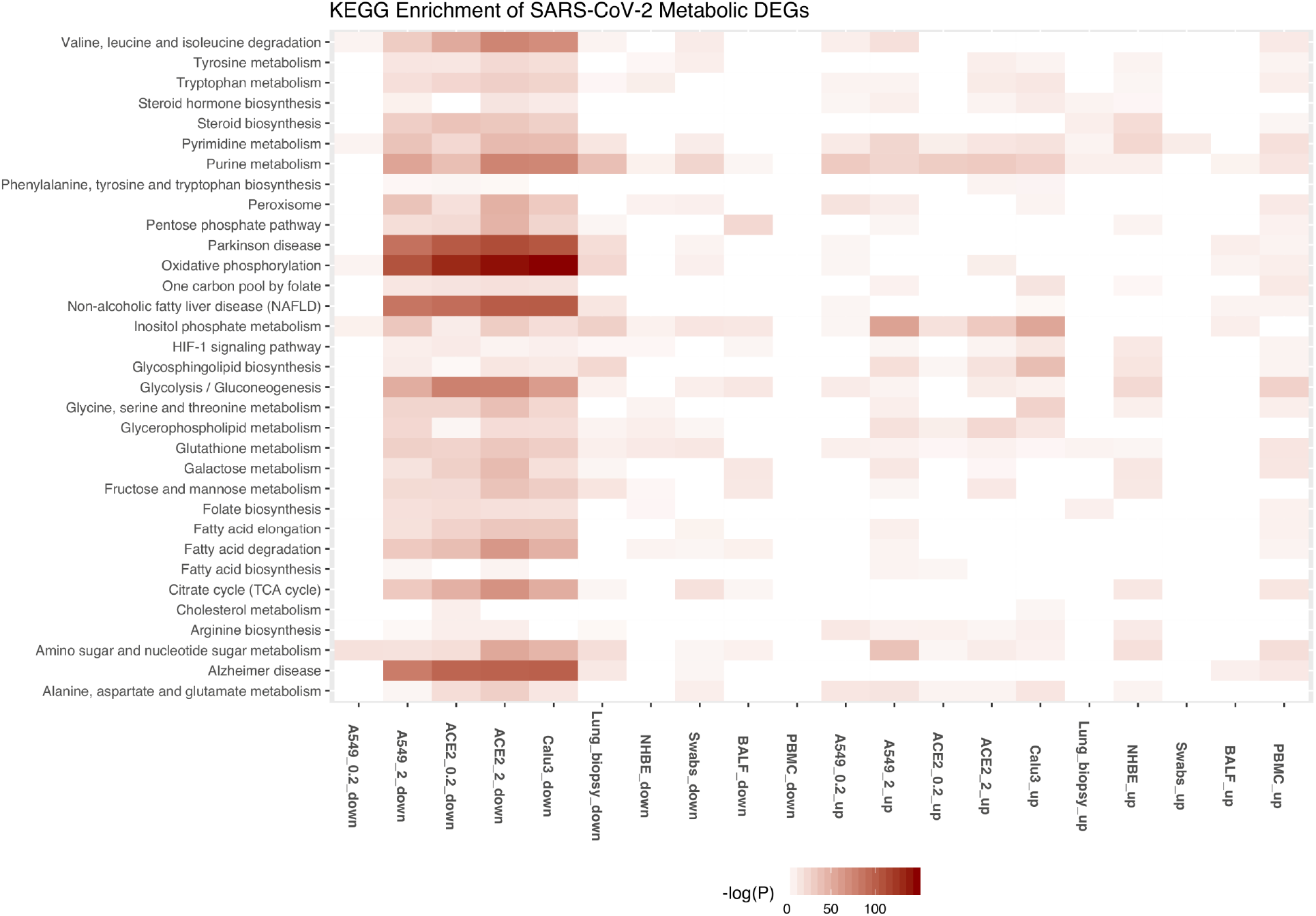
KEGG pathways associated with upregulated and downregulated metabolic genes in different cell lines and patient samples with SARS-CoV-2 infection. A549 and A549-ACE2 is shown for MOI of 0.2 (A549_0.2, ACE2_0.2) and 2 (A549_2, ACE2_2). Up and down represent the upregulated and downregulated metabolic DEGs.

**Figure 3:**
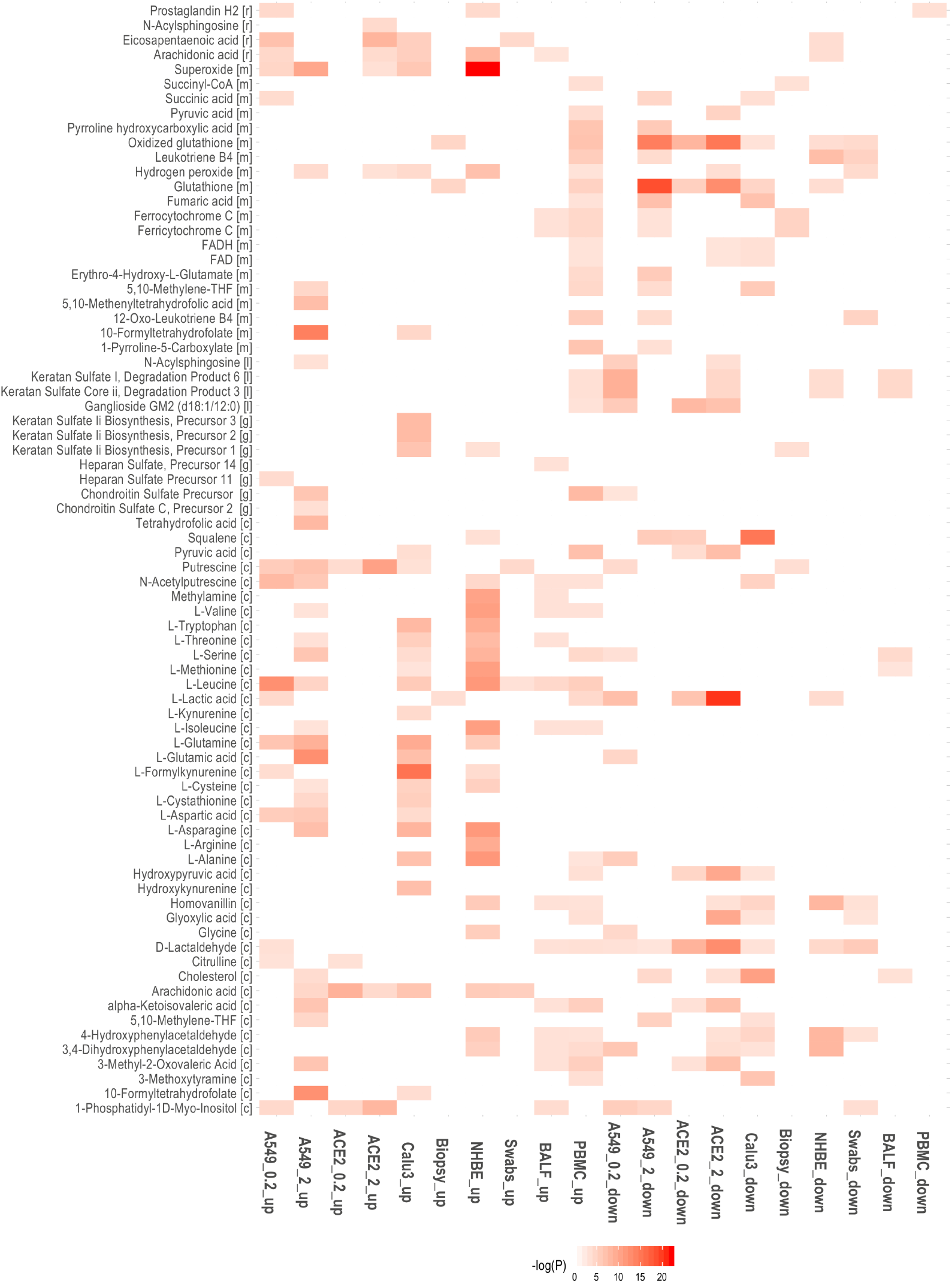
Downregulated and upregulated reporter metabolites in different cell lines and patient samples with SARS-CoV-2 infection. A549 and A549-ACE2 is shown for MOI of 0.2 (A549_0.2, ACE2_0.2) and 2 (A549_2, ACE2_2). Up and down represent the upregulated and downregulated metabolic DEGs.

Interestingly, we observed that lactic acid is also a reporter metabolite in multiple conditions, including PBMC. Lactate dehydrogenase (LDHA) gene is significantly upregulated in PBMC (**Table S2**). This resembles a condition of aerobic glycolysis termed as “Warburg effect”, which is a hallmark of cancer. The Warburg effect is also reported with oncogenic viruses’ infection. Lactic acid is shown to influence the cytokine production (Haas et al., 2015; Ratter et al., 2018). The activation of PI3K-Akt-mTOR pathway controls the Warburg effect in cancers and viral infection (Mannova and Beretta, 2005; Martin et al., 2012; Noch and Khalili, 2012; Su et al., 2014). PI3K controlled inositol phosphate metabolism is upregulated in Calu3 and A549 with MOI=2 (A549_2).

Further, lactate is also shown to be a natural suppressor of antiviral signaling. Studies show that lactate inhibits RLR signaling by binding to the MAV transmembrane domain (Ren et al., 2020; Zhang et al., 2019). LDHA inhibition protects the mice from viral infection by increasing the interferons production. Therefore, pharmacologically targeting LDHA is a promising strategy to induce antiviral signaling. RLR signaling also is shown to suppress glycolysis by inhibiting the enzyme hexokinase HK2 (Zhang et al., 2019). This mutual antagonism between RLR and glycolysis may play a role in SARS-Cov-2 infection (Stoolman and Chandel, 2019). We observed that glycolytic enzymes are upregulated in only NHBE (HK2, ENO1) and PBMC (ENO1, GAPDH, TPI1, PKM, PGM2, HK1) (**Table S2**). Glycolytic genes TPI1, ENO1, PKM, PGAM1, GAPDH, and PFKL are downregulated in A549_2, A549_ACE2 with MOI=0.2 and 2 (ACE2_0.2, ACE2_2), and Calu3. However, SLC2A1 (GLUT1) and HK2 are upregulated in these cell lines. HK2 integrates both energy metabolism and cell protection via Akt/mTOR signaling pathway (Roberts and Miyamoto, 2015). It is shown to enhance autophagy by inhibiting the mTOR activation. Further, the precursors of chondroitin sulphate and keratan sulphate (glycosaminoglycans) synthesis are also reporter metabolites in A549_2 and Calu3 while the degradation products of keratan (in lysosome) is reporter metabolites in PBMC (**Figure 3**).

Genes involved in the conversion of serine (SDS and SDSL) and alanine (GPT2) to pyruvate are upregulated in Calu3 (**Table S2**). Serine is a reporter metabolite in most infected conditions. We also found that the HIF1A pathway is enriched in SARS-CoV-2 transcriptome (**Figure 2**). Some of its targets, such as VEGF, are upregulated in A549_2, ACE2_0.2, ACE2_2, Calu3 and NHBE (**Table S1**). On the other hand, we found that VHL that targets HIF1A to degradation are also upregulated in these cell lines. HIF1A promotes a shift in mitochondrial metabolism to anaerobic glycolysis and is also regulated by mTOR (Dodd et al., 2015). These results suggest a dysfunction in the PI3K-Akt-mTOR-HIF1A axis in SARS-Cov-2. We propose that PI3K inhibitors Wortmannin and LY294002, which are previously known to have anti-cancer activity and inhibit Warburg effect, may be repurposed for the antiviral therapy. These inhibitors are shown to inhibit the Middle East Respiratory Syndrome coronavirus (MERS-CoV) replication *in vitro* (Kindrachuk et al., 2015).

### Redox homeostasis

We also found that genes of the pentose phosphate pathway (PPP) and folate metabolism (DHFR, SHMT1, MTHFD1) are downregulated in some cell lines (**Table S2**). These pathways play an important role in maintaining the redox balance by the production of NADPH. Further, we observed that the *de novo* synthesis of glutathione is also downregulated (**Figure 2**). These may reflect the oxidative stress condition of the host. Glutathione deficiency is also considered to be a severe manifestation of SARS-Cov-2 (Polonikov, 2020). However, we found that SOD2 involved in the conversion of reactive oxygen species (ROS) to peroxides is upregulated in most cell lines. Further, NADPH oxidase enzyme 2 (DUOX2) is upregulated in most cell lines (**Table 2**). They are usually expressed in epithelial tissues lining the respiratory and intestinal tracts and participate in the host defense against microbial infection at mucosal surfaces by the generation of peroxides (Geiszt et al., 2003; Joo et al., 2012). It is also an antiviral gene induced in response to cytokines (Fink et al., 2013; Grandvaux et al., 2015). In PBMC, genes of folate metabolism (SHMT1, MTHFD2) are upregulated along with CAT, PRDX, and GPX1 involved in the elimination of peroxides (**Table S2**). MTHFD2 is also upregulated in A549_2, Calu3 and NHBE. The metabolites of homocysteine metabolism (L-cysteine and Cystathionine) are significantly upregulated in A549_2 and Calu3.

### Polyamine metabolism

Genes that control polyamine levels are also altered in some cell lines. This includes ODC1 and SAT1, which are known to influence the polyamines’ synthesis and catabolism, respectively. ODC1 is involved in the generation of putrescine, and SAT1 is involved in catabolizing spermine back to spermidine and putrescine. We found putrescine, N-acetylputrescine, and L-arginine as reporter metabolites of infected conditions. Polyamines are positively charged and bind DNA/RNA to assist in viral replication (Mounce et al., 2017). They help to neutralize the negative charges of DNA and assist in packing of DNA into viral particles(Lanzer and Holowczak, 1975; Lightfoot and Hall, 2014). Studies show that MERS-CoV is dependent on polyamines for replication (Mounce et al., 2017). Polyamines are known to control Myc expression, and Myc, in turn, controls the transcription of genes ODC1, SRM and SMS involved in polyamine synthesis (positive feedback loop) (Firpo and Mounce, 2020) (**Figure 4**). Therefore, the depletion of polyamines can be a strategy to reduce viral infection.

**Figure 4:**
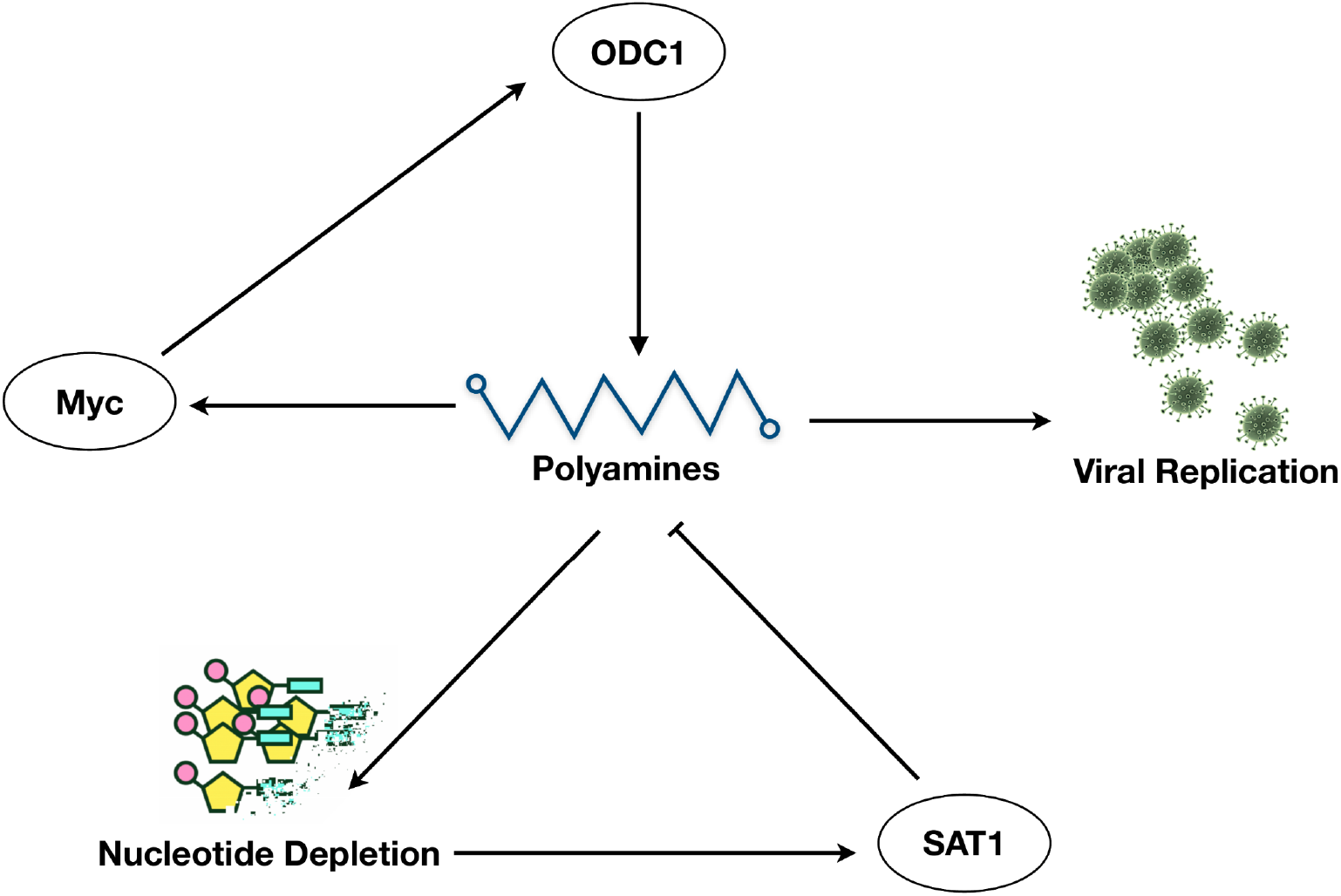
Feedback loop regulation of polyamine synthesis controls viral replication.

ODC1 inhibitor Difluoromethylornithine (DFMO), which is in clinical trials for cancer treatment, can be tested for antiviral effect against SARS-Cov-2(Mounce et al., 2016; Raniga and Liang, 2018). Further, (FDA)-approved antiviral ribavirin enhances SAT1, which depletes polyamines levels and viral titers (Tate et al., 2019). Polyamine levels are regulated by a negative feedback loop involving SAT1 (**Figure 4**). Ribavirin is proposed as a novel coronavirus treatment strategy (Khalili et al., 2020). Ribavirin is also part of triple-antiviral therapy proposed for alleviating symptoms in mild to moderate SARS-CoV-2 conditions (Hung et al., 2020). Further, purine and pyrimidine metabolism are also significantly downregulated in most conditions (**Figure 2**). However, we observe CMPK2, NT5E, and TYMP involved in the pyrimidine metabolism are upregulated in different conditions (**Table S2**). CMPK2 is an interferon-stimulated gene known to restrict HIV infection (El-Diwany et al., 2018). CMPK2 is found adjacent to gene RSAD2, which is well known to inhibit a wide variety of RNA and DNA viruses by synthesizing chain terminators that block DNA/RNA synthesis (Gizzi et al., 2018).

### Aromatic, Arginine, and branched chain amino acid metabolism

Genes involved in Tryptophan catabolism (KYNU, KMO, IDO1) are upregulated in multiple cell lines along with the upregulation of nicotinamide phosphoribosyltransferase (NAMPT) suggesting that the *de novo* synthesis of NAD from tryptophan may be active (**Table 2 and Table S2**). We found L-formylkynurenine and kynurenine of tryptophan metabolism as reporter metabolites. IDO1-mediated tryptophan depletion in the short term is shown to inhibit the viral replication but leads to immunosuppression in the long term (Raniga and Liang, 2018). Further, metabolites of tyrosine metabolism (homovanillin and 3-methoxytyramine) are upregulated in PBMC (**Figure 3**).

We found that argininosuccinate synthetase 1 (ASS1) involved in *de novo* arginine synthesis from aspartate is upregulated in most cell lines (**Table S2**). Inhibition of ASS1 is shown to increase viral replication, and its deficiency affects the immune cell activation(Grady et al., 2013; Tarasenko et al., 2015). This suggests that ASS1 is part of the antiviral pathway involved in the host defense. ASNS, which is involved in the synthesis of asparaginase from aspartate, is upregulated in A549 and Calu3. Aspartic acid, asparaginase, and citrulline are also reporter metabolites of infected conditions (**Figure 3**). Asparaginase is found to be the novel host factor promoting the replication of human cytomegalovirus (HCMV) (Lee et al., 2019). Valine, leucine, and isoleucine (branched chain amino acids, BCAAs) degradation pathway is also downregulated in some cell lines (**Figure 2**). The degradation of BCAAs generates CoA products that feed into the TCA cycle. L-isoleucine, L-leucine, L-valine, and alpha-ketoisovaleric acid are reporter metabolites of infected conditions (**Figure 3**). BCAAs are reported to activate mTOR and the interferon signalling by inactivating the suppressor of cytokines signaling 3 (SOCS3) (Honda et al., 2011). BCAA inhibits hepatitis C virus (HCV) replication and promotes infectious particle formation (Ishida et al., 2013). In PBMC, genes involved in BCAA degradation are upregulated. Further, BCAT1 is upregulated in A549_2 and Calu3. The first step of BCAA degradation (BCAT1 and BCAT2) can induce metabolic reprogramming to combat the ROS (Wang et al., 2019).

### Lipid metabolism

Genes of fatty acid degradation and elongation are downregulated in most cell lines except for PBMC (**Table S2**). FASN that is involved in fatty acid synthesis is also downregulated. However, the gene involved in the hydrolysis of triacylglycerol (LPL) is upregulated. Further, HMGCS1 and HMGCR involved in cholesterol synthesis from acetyl CoA are downregulated (A549_2, ACE2_0.2, ACE2_2, Calu3) while steroid hormone synthesis pathway from cholesterol is upregulated. Interestingly, CH25H, which is involved in converting cholesterol to cholesterol 25-hydroxylase (25HC), is upregulated in ACE2_2 and Calu3. This is an interferon regulated gene, and 25HC is known to have an antiviral effect on HIV, nipah virus, and ebola by blocking the membrane fusion of the virus (Liu et al., 2013). The distribution of cholesterol is shown to be important for viral replication, entry, and budding (Raniga and Liang, 2018). The cholesterol and fatty acid synthesis are regulated by transcription factor SREBPs, which is activated by a viral infection (HCV) through PI3K/Akt pathway (Jackel-Cram et al., 2010). Gene encoding LDL receptor (LDLR) is also upregulated, which RNA viruses use for the entry into the host (Finkelshtein et al., 2013; Molina et al., 2007).

SGMS1 and SGMS2 involved in *de novo* synthesis of sphingolipids are upregulated in A549_2, ACE2_2, and Calu3 (**Table S2**). In A549_2, serine palmitoyltransferase (SPTLC1, SPTLC2) and ceramide synthase (CERS2, CERS5, CERS6) are also upregulated. Further, genes of glycosphingolipid synthesis (B4GALT1, B3GNT5, B3GNT2) and glycerophospholipid metabolism (PLA2G4A, PLA2G4C, PLD1, PLD2) are also upregulated. Phospholipase 2 hydrolyzes membrane phospholipids to release arachidonic acid. PTGS2 involved in the conversion of arachidonic acid to prostaglandin is also upregulated. Both arachidonic acid and prostaglandin H2 are reporter metabolites in the infected conditions (**Figure 3**). HCV and dengue virus depend on MAPK activated PLA2G4C that helps in targeting core protein to lipid droplets (Menzel et al., 2012).

### Methytranferases and Heme catabolism

We found SETD2 and ASH1L that encode histone methyltransferases to be upregulated in A549_2, ACE2_0.2, ACE2_2, and Calu3. Methylation of STAT1 by SETD2 is shown to be critical for interferon antiviral activity (Chen et al., 2017). Further, ASH1L also plays a role in the suppression of TLR-mediated immune response (Xia et al., 2013). HMOX1 and HMOX2, which are known to have an antiviral effect, are downregulated (Soota and Maliakkal, 2014; Zhu et al., 2017) (**Table S2**). HMOX1 protects cells from programmed cell death by catabolizing free heme and its induction can have a therapeutic effect in different immune-mediated inflammatory diseases (Gozzelino et al., 2010). HMOX1 knockout mice showed significantly higher cytokine responses (Soota and Maliakkal, 2014). It is required for the protection against inflammatory insult to the lung.

### Host-pathogen interactome and transcriptional regulation of metabolic genes

The host-pathogen interactome of SAR-CoV-2 was used to identify the direct and indirect metabolic targets of viral proteins (**Figure 5 and Figure S2**). We found that viral proteins target the host mitochondrial metabolism (M^Pro^, Nsp7, Orf9c), purine and pyrimidine synthesis (Nsp1, Nsp14), glutathione metabolism and oxidative stress (Nsp5_C145A, Orf3a), lipid metabolism (M^Pro^, Nsp2, Nsp7, Orf10), folate metabolism (N, Orf8), chondroitin biosynthesis (Orf8) and prostaglandin biosynthesis (Nsp7). SARS-CoV-2 proteins also target different metabolic genes through host protein kinase A (Nsp13), casein kinase 2 (N), G-protein subunits (Nsp7), Histone deacetylase 2 (Nsp5), Catechol-O-methyltransferase (Nsp7), Galactosidase Alpha (Nsp14) and NF-Kappa-B-activating kinase (Nsp13). G-protein subunits GNG5 and GNB1 control the activation of PI3K/Akt pathway by directly targeting the PI3K subunits.

**Figure 5:**
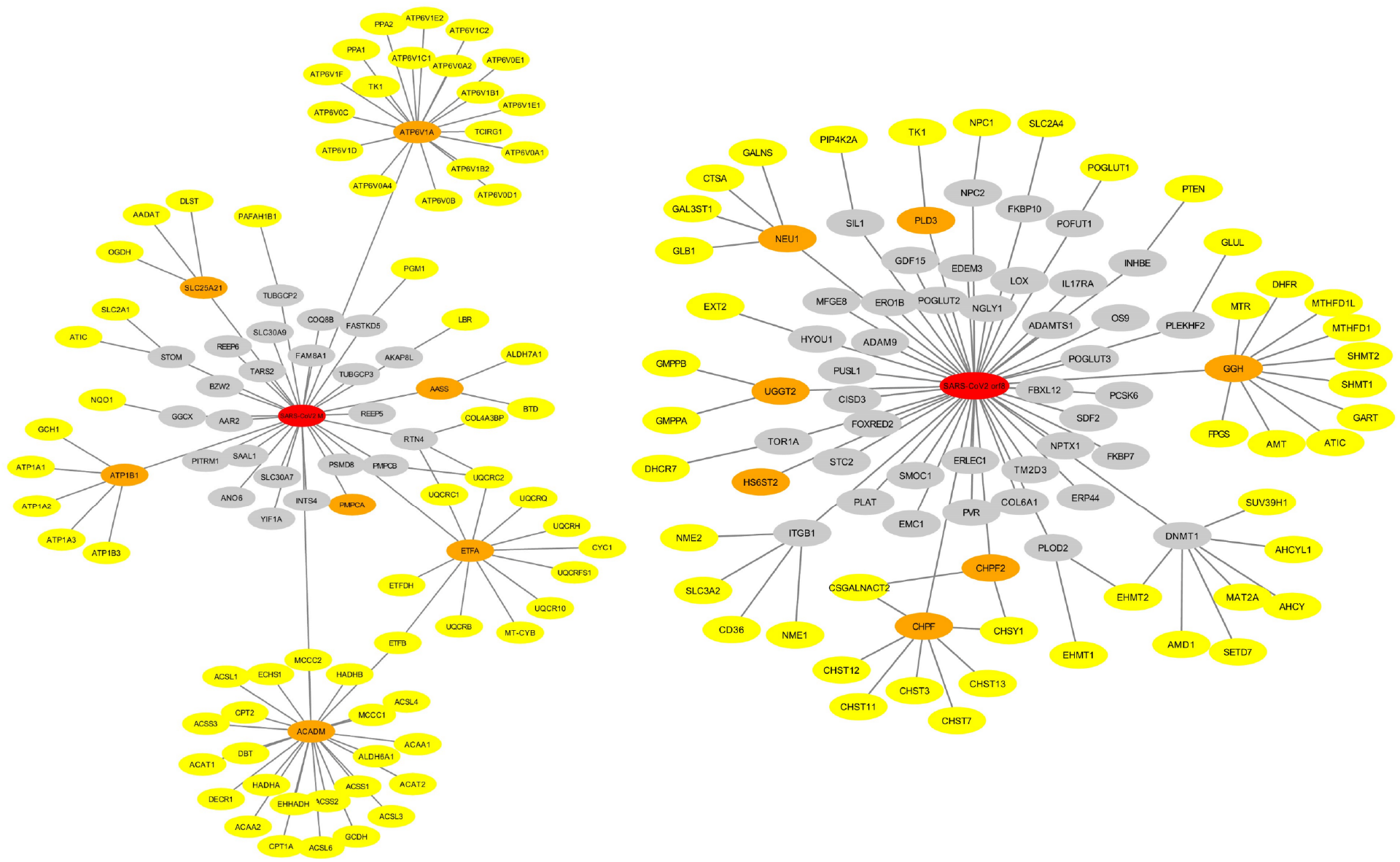
Protein-protein interaction of viral proteins M^Pro^ (left) and Orf8 (right) with the host metabolic proteins. Viral proteins are shown in red color. Metabolic DEGs interacting directly and indirectly with viral proteins are shown in orange and yellow colors, respectively. Non-metabolic genes are shown in the grey color.

To understand the transcriptional regulation of metabolism, we identified the potential transcription factors based on the binding site enrichment of DEGs under each condition. We found an overlapping set of transcription factors is associated with upregulated and downregulated genes in different conditions (**Tables 3 and 4**). This includes transcription factors of NFκB family (RELA, NFKB1) that are known to control the transcription of host immune response. NFκB is activated by different viruses, including HCV, hepatitis B virus (HBV), epstein-barr virus (EBV), human immunodeficiency virus (HIV-1), and influenza virus (Hiscott et al., 2001), which helps in viral replication and suppressing host cell death. Transcription factors HIF1A, NFE2L2/NRF2, PPARA, and SERBF1, are also associated with differentially expressed genes. NFE2L2 is involved in the oxidative stress, while SERBF1 and PPARA are involved in lipid homeostasis (Brown and Goldstein, 1997; Kobayashi and Yamamoto, 2006). PPARA stimulation by DNA virus infection suppresses the interferon signaling and impairs immunity against the virus (Tao et al., 2019). PPAR controls peroxisome metabolism, which is shown to be important for different viruses. We observed that the peroxisome function is altered in different cell lines and human samples (**Figure 2**).

**Table 3:** Transcription factor (TF) significantly (p < 0.05) associated with upregulated metabolic DEGs. Number of occurrences of each TF in different host conditions are given.

| TF | Occurrences | Samples |
| --- | --- | --- |
| SP1 | 9 | A549-0.2, A549-2, ACE2-0.2, ACE2-2, Calu3, Lung_Biopsy, NHBE, BALF, PBMC |
| HIF1A | 8 | A549-0.2, A549-2, ACE2-0.2, ACE2-2, Calu3, NHBE, BALF, PBMC |
| USF1 | 8 | A549-0.2, A549-2, ACE2-0.2, ACE2-2, Calu3, Lung_Biopsy, BALF, PBMC |
| NFKB1 | 8 | A549-0.2, A549-2, ACE2-0.2, ACE2-2, Calu3, Lung_Biopsy, NHBE, Swabs |
| RELA | 8 | A549-0.2, A549-2, ACE2-0.2, ACE2-2, Calu3, Lung_Biopsy, NHBE, Swabs |
| PPARA | 7 | A549-0.2, A549-2, ACE2-0.2, ACE2-2, Calu3, Lung_Biopsy, NHBE |
| USF2 | 7 | A549-0.2, ACE2-0.2, ACE2-2, Calu3, Lung_Biopsy, BALF, Swabs |
| PPARG | 7 | A549-0.2, A549-2, ACE2-0.2, ACE2-2, Calu3, Lung_Biopsy, NHBE |
| NR0B2 | 7 | A549-0.2, A549-2, ACE2-0.2, ACE2-2, Calu3, NHBE, PBMC |

**Table 4:**
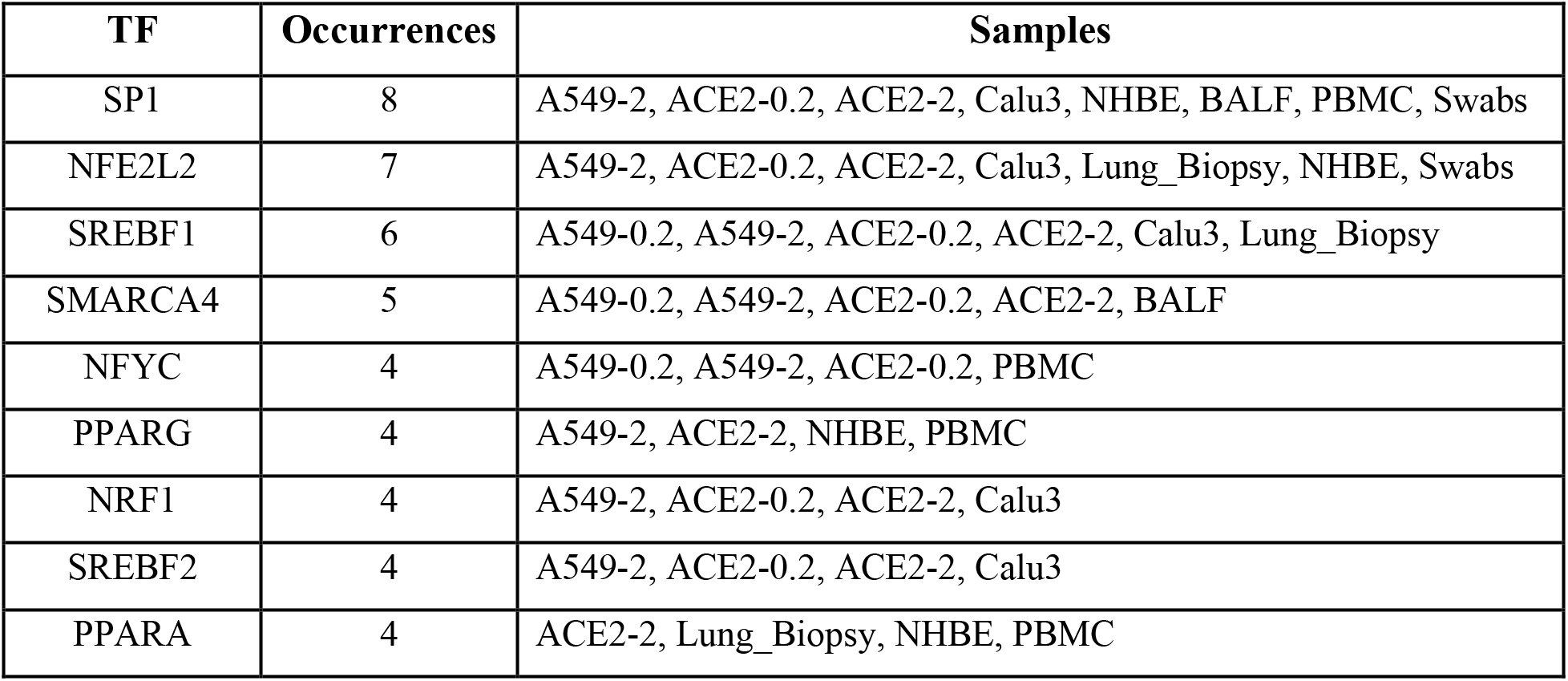
Transcription factor (TF) significantly (p < 0.05) associated with downregulated metabolic DEGs. Number of occurrences of each TF in different host conditions are given.

### Conclusions

In summary, our study performs a comprehensive analysis of several transcriptomic data and generates hypotheses into how the host metabolic pathway is hijacked in response to SARS-CoV-2. We observed different metabolic changes that are pro- and antiviral in nature. A shift in this balance may play a role in disease pathogenesis. We provide a mechanistic view of how the antiviral metabolism can be targeted/enhanced for reducing viral titers. We suggest probable targets in carbohydrate and amino acid metabolism, polyamine synthesis and redox homeostasis. These warrant further exploration with more samples and in vitro studies to test predictions.

## Supplementary information

**Table S1:** Differentially expressed genes in response to SARS-CoV-2 in different cell lines and patient samples.

**Table S2**: Differentially expressed metabolic genes in response to SARS-CoV-2 in different cell lines and patient samples.

**Figure S1**: Overlap of differentially expressed genes in different cell lines and patient samples with SARS-CoV-2 infection.

**Figure S2**: Protein-protein interaction of viral proteins with the host metabolic proteins. Viral proteins are shown in red color. Metabolic DEGs interacting directly and indirectly with viral proteins are shown in orange and yellow colors, respectively. Non-metabolic genes are shown in the grey color.

